# Intrinsic disorder in flaviviral capsid proteins and their role in pathogenesis

**DOI:** 10.1101/2023.09.26.559664

**Authors:** Anirudh Sundar, Pavithra Umashankar, Priyanka Sankar, Kavitha Ramasamy, Sangita Venkataraman

## Abstract

A high level of disorder in many viral proteins is a direct consequence of their small genomes, which makes interaction with multiple binding partners a necessity for infection and pathogenicity. A segment of the flaviviral capsid protein, also known as the molecular recognition feature (MoRF), undergoes a disorder-to-order transition upon binding to several protein partners. In order to understand their role in pathogenesis, the MORFs were identified and their homology was studied. Despite the lack of sequence similarities, the docking studies of Cs with the host proteins indicate conserved interactions involving MORFs across members of the phylogenetic subclades. Additionally, it was observed from the protein-protein networks that some MoRFs preferentially bind proteins that are involved in specialized functions such as ribosome biogenesis. The findings point to the importance of the MoRFs in the flaviviral life cycle, with important consequences for disease progression and suppression of the host immune system. Potentially, they might have impacted the way flaviviruses evolved to infect varied hosts using multiple vectors.

## 1. Introduction

The lack of a robust three-dimensional structure in many proteins is invaluable to their cellular functions (including but not confined to signaling, recognition, and regulation), enhancing specificity and low-affinity interactions with multiple partners (Mishra et al. 2020). Over 10% of the total entries across all organisms in the DisProt database (Quaglia et al. 2022), which contains curated disordered proteins, belong to viruses (Wubben et al. 2020). Previous studies have shown that Intrinsically Disordered Regions (IDRs) are at the core of important protein-ligand interactions in flaviviruses (Mishra et al. 2020; Martins and Santos 2020). The IDRs offer high flexibility to viral proteins and help them quickly adapt to a changing environment and evade the defense mechanisms of the host, potentially by hindering recognition by components of the immune system, and thus are correlated with virulence (Goh et al. 2016, 2019).

The genus *flavivirus* belonging to the family *flaviviridae* consists of multiple *arboviruses* like *Dengue virus* (DENV), *Zika virus* (ZIKV), *West Nile virus* (WNV), and *Japanese encephalitis virus* (JEV) that are the cause of life-threatening neurotropic, visceral, and congenital diseases in humans. These viruses possess positive-sense RNA genomes that encode three structural proteins (C, envelope-E, and precursor membrane protein-prM) and seven non-structural proteins (NS1, NS2A, NS2B, NS3, NS4A, NS4B, and NS5). The C engages in vital processes like entry, translational shutoff, and transcriptional regulation that are integral to infection and interacts with many cellular proteins, including RNA-binding proteins such as La, cellular nucleoporins, and importin α/β (Shah et al. 2018; Selinger et al. 2022). For instance, prior research has shown the ability of specific DENV2 C domains to cross the blood-brain barrier through adsorptive-mediated transport (AMT) (Neves et al. 2017). Several host proteins involved in RNA processing and mitochondrial ribosomes have been shown to interact with DENV C, while ZIKV C is thought to interact with proteins involved in transcription, DNA replication, and proliferation (Shah et al. 2018). Examination of similarities in the C proteins at a structural level and phylogenetic comparison of their sequences indicate their potential to be good evolutionary markers (Faustino et al. 2019). Flaviviral Cs also exhibit RNA binding properties at the N and C terminal MoRFs, with high affinity and low specificity, regardless of a definitive 3-D structure (Ivanyi-Nagy et al. 2008; Byk and Gamarnik 2016).

Given the importance of disordered MoRFs in non-specific interactions, we identified the MoRFs and investigated the patterns of disorder in flaviviral Cs. Additionally, we examined the composition, distribution, and homology of MoRFs across the members of *Flaviviridae.* We also investigated the role of MoRFs in binding to various host proteins using a systems biology approach and analyzed how such relationships evolved. Analysis of viral-host protein-protein interactions (PPI) mediated via the MoRFs provides insight into the multifunctionality of the Cs and may contribute to the development of new antiviral therapies.

## 2. Materials and methods

### 2.1 Multiple sequence alignment and phylogenetic analysis

The flaviviral genome polyprotein sequences were collected from the SwissProt database (Bateman et al. 2021). Out of the 124 sequences, a total of 25 genome polyproteins were utilized for further analysis (UniProt IDs: P05769, Q91B85, Q01299, P09732, P29837, P22338, Q04538, Q7T6D2, Q5WPU5, Q32ZD7, Q32ZD5, D7RF80, C8XPA8, C5H431, C8XPB2, Q32ZD4, Q32ZE0, Q32ZE1, Q074N0, Q9Q6P4, G3FEX6, P33478, P29990, P27915, Q58HT7) (**Table 1**). The Cs were annotated using the Pfam (Paysan-Lafosse et al. 2023) option in IUPred3 (Abor Erd et al. 2021) using the long disorder option and were cataloged for further analysis (**Supplementary Table S1**). The sequences were aligned with MUSCLE, and subjected to phylogenetic analysis using the maximum likelihood method with 1000 bootstraps in MEGA11 (Tamura et al. 2021) (**Fig 1**).

**Fig 1:**
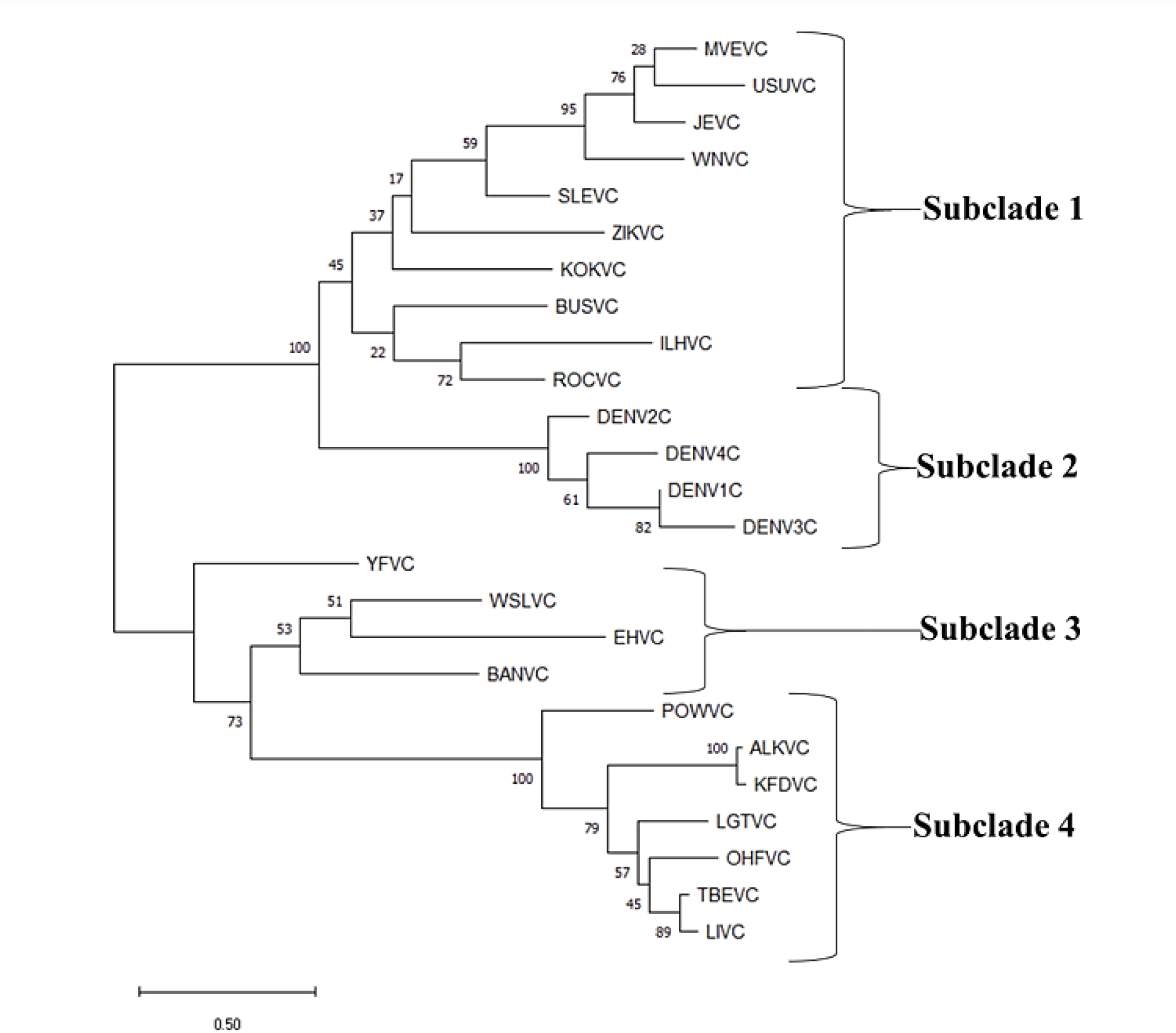
Phylogenetic tree. Maximum Likelihood rooted tree constructed for 25 Capsid sequences of the flaviviruses using MEGA11. The confidence was assessed with 1000 bootstrap replicates. (Tamura et al. 2021)

**Table 1:**
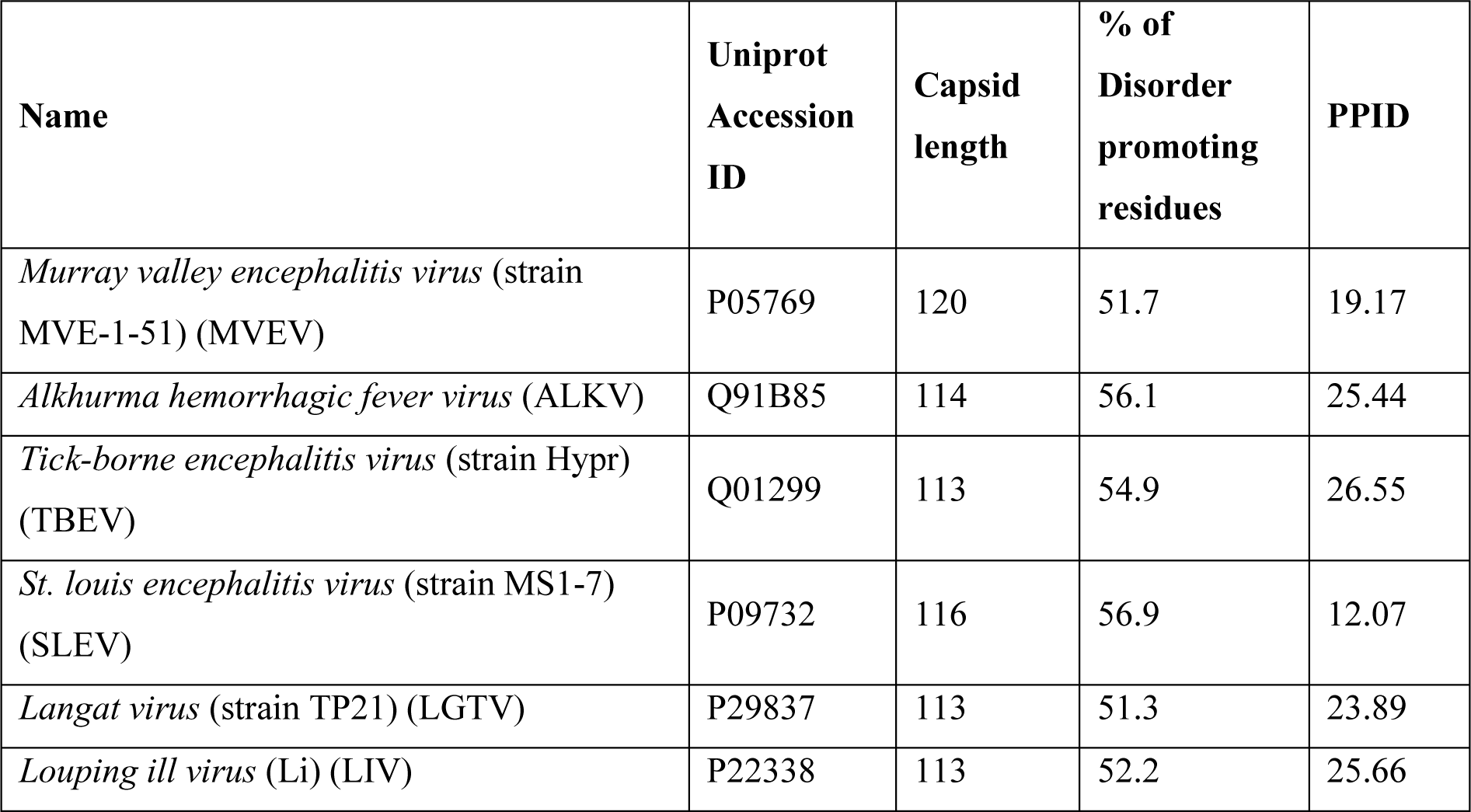
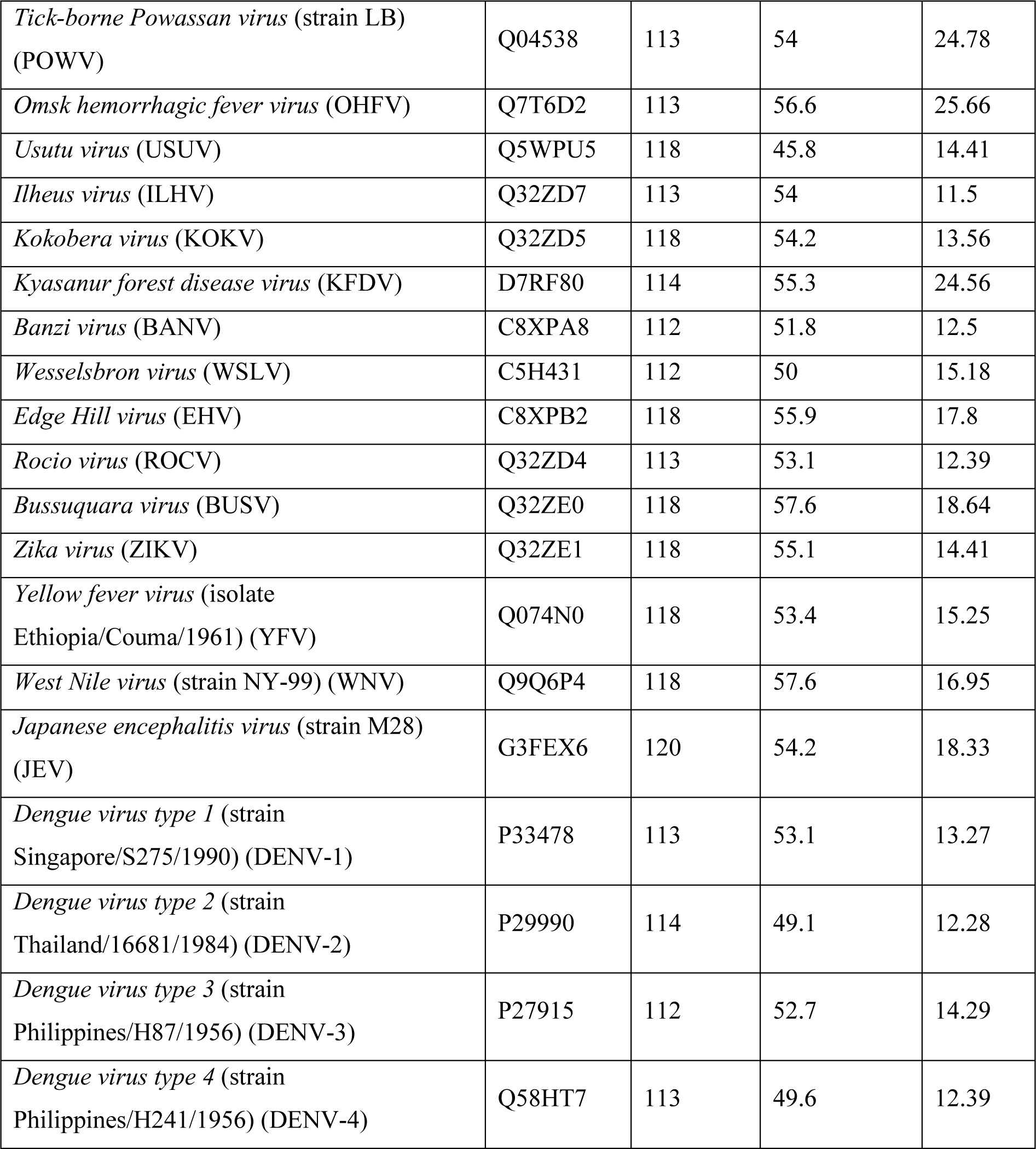
Details of viral capsid proteins. Uniprot Ids of retrieved polyprotein sequences, length of capsid sequences identified using Pfam, percentage of disorder promoting residues in the capsid, and Predicted Percentage of Intrinsic Disorder (PPID) in the capsid. (Bateman et al. 2021; Abor Erd et al. 2021; Paysan-Lafosse et al. 2023)

### 2.2 Disorder analysis of Cs

Composition profiling was carried out to identify the proportion of order- and disorder-promoting residues in the Cs (**Supplementary Table S2)**. Since amino acids like Ala, Arg, Gly, Gln, Ser, Glu, Asp, His, Lys, and Pro are prominent in IDRs, they are considered to be disorder promoting (**Table 1**) (Redwan et al. 2019). The overall percentage of the disorder promoting residues were computed for the individual Cs and their compositional bias was studied.

The disordered regions in the Cs were predicted using PrDOS (Ishida and Kinoshita 2007). Residues having a disorder probability greater than 0.5 were considered disordered. The data thus obtained were used to generate disorder probability plots (**Fig. 2**). Additionally, the PPID (Predicted Percentage of Intrinsic Disorder) (Sharma et al. 2021) was calculated for the C sequences (**Table 1**) using the formula (Barik 2020):

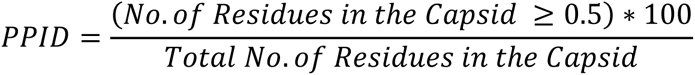

**Fig 2:**
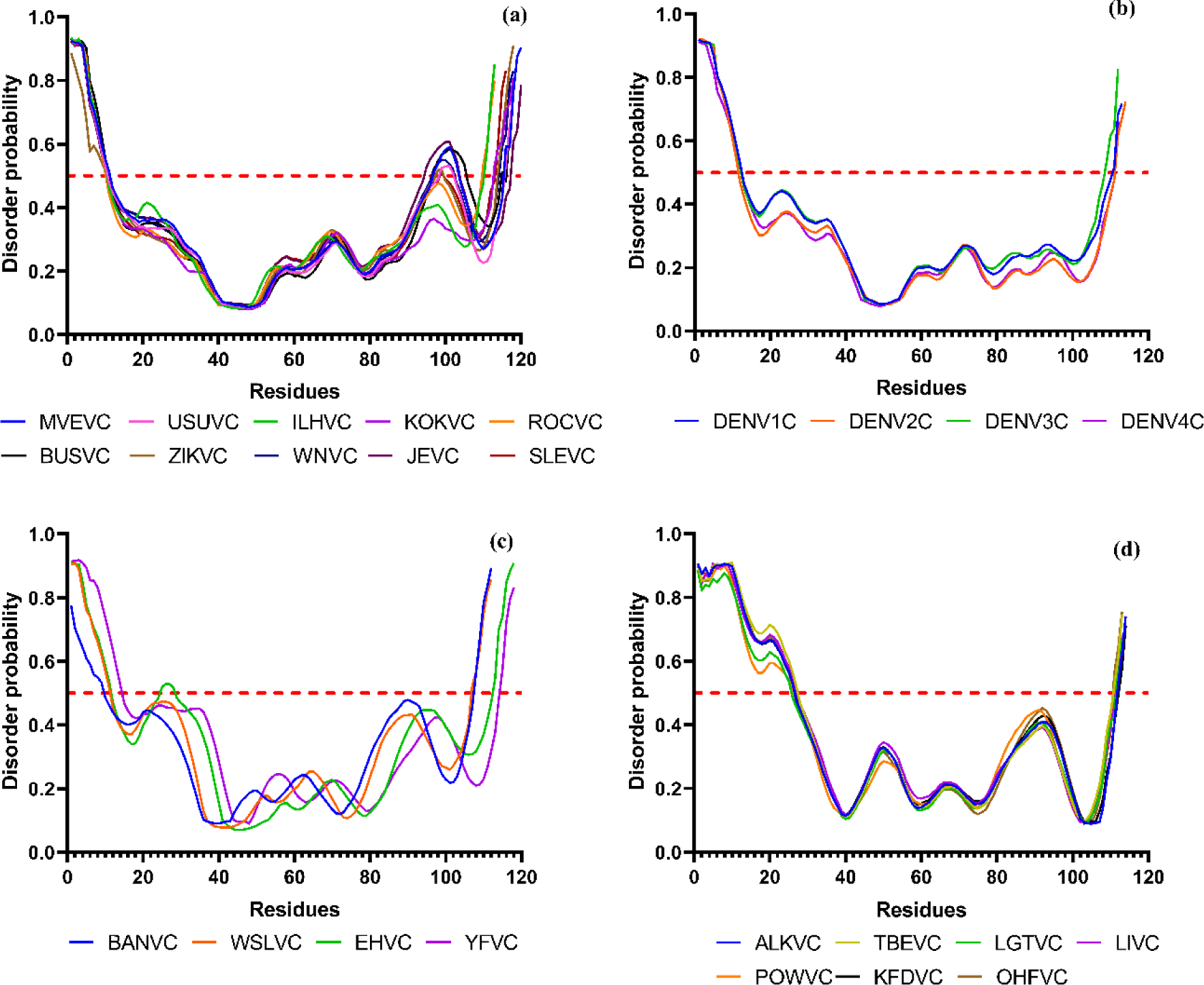
Combined Disorder Probability Plots. Disorder probability for Capsids plotted using scores obtained from PrDOS (a) Capsid sequences from subclade 1, (b) Capsid sequences from subclade 2, (c) Capsid sequences from YFV and subclade 3, and (d) Capsid sequences from subclade 4 are shown. (Ishida and Kinoshita 2007)

Web-based predictors MoRFCHiBi_Web (MCW) (Malhis et al. 2016) and DISOPRED3 (Jones and Cozzetto 2015) were used to predict protein binding regions that are unique to IDRs (MoRFs). Both MCW and DISOPRED3 use SVM-based models for the prediction of MoRFs. While the former calculates six propensity scores based on local physicochemical properties of the amino acids to predict MoRFs, the latter combines two independent predictors of disorder (a neural network and a nearest neighbor classifier) to identify disordered regions with MoRFs.

### 2.3 Interactome analysis and functional enrichment studies

The interactomes for selected viruses were obtained from the literature (Shah et al. 2018; Li et al. 2019; Selinger et al. 2022) and C-specific interactions were filtered (**Supplementary Table S3)**. PPI networks for the Cs of individual viruses were constructed using Cytoscape 3.9.1 (Shannon et al. 2003) and the genes encoding common interacting partners were highlighted. Functional enrichment was carried out using gProfiler (Raudvere et al. 2019) to discover GO categories that were enriched for human proteins (**Supplementary Table S5**).

### 2.4 Molecular docking

Molecular docking was performed using HDOCK to analyze the mode of binding of the Cs with the 14 common interacting host proteins (Yan et al. 2017, 2020). The structures of dimeric Cs were retrieved from the four available PDB entries (TBEV C: 7YWQ, DENV2 C: 1R6R, ZIKV C: 6C44, and WNV C: 1SFK). For the human proteins, PDB/AlphaFold structures were used depending on the availability (TSR1: 6G18, DDX18: 3LY5, GNL3: AF-Q9BVP2-F1, RRP1B: AF-Q14684-F1, RSL1D1: AF-O76021-F1, SRPK1: 1WAK, ZC3HAV1: AF-Q7Z2W4, YTHDC2: AF-Q9H6S0-F1, DKC1: 7TRC, PURA: AF-Q00577-F1, SRSF5: AF-Q13243-F1, SSB: AF-P05455-F1, XRCC6: 1JEY, NOP16: AF-Q9Y3C1-F1) (Berman 2000; Jumper et al. 2021; Varadi et al. 2022). The interacting residues were obtained from the complex with the highest docking score in HDOCK and analyzed by comparing them to the MoRFs obtained from MCW and DISOPRED3. The percentage of MoRF residues found to interact with the host proteins was computed for each docked complex (**Supplementary Table S4**). Using the docking results, a Venn diagram was generated using the web tool, http://bioinformatics.psb.ugent.be/webtools/Venn/ (Yves Van de Peer) to represent the number of common interacting proteins between the four viruses. The percentage of MoRF residues interacting with the common host partners was calculated for individual C to elucidate the role of IDRs in protein binding. A heatmap was then constructed using R studio (R 4.2.2) (RStudio Team) to visualize and interpret the data.

## 3. Results

### 3.1 Multiple sequence alignment and phylogenetic analysis

Multiple sequence alignment was done for the 25 flavivirus C sequences (**Supplementary Table S1**), and a phylogenetic tree was constructed by the maximum likelihood method using MEGA11 (Tamura et al. 2021) (**Fig 1**). The consensus tree following bootstrapping showed four subclades, with YFV as the outgroup. The flaviviruses were distributed in two main clades, with all the tick-borne viruses in subclade 4. A cross-neutralization study of flaviviruses using their corresponding antisera, performed in 1974 (De Madrid and Porterfield 1974), classified them as:

- Subgroup 1 - LGTV, KFDV, LIV, OHFV, TBEV
- Subgroup 3 - JEV, MVEVC, WNV, SLEV, USUV, KOCV
- Subgroup 4 - ZIKV
- Subgroup 6 - BANV, EHV
- Subgroup 7 - DENV1, DENV2, DENV3, DENV4
- Outliers - BUSV, WSLV, ILHV, POWV, YFV

Thus, the subclades observed in the phylogenetic tree were corroborated by those identified in the cross-neutralization study. The majority of the branches had bootstrap support of 60% and above. From the MSA (**Supplementary Fig S1**) it is evident that certain sequences were conserved within tick-borne and mosquito-borne viruses, respectively. The 10N-M-L-K-R14 motif of capsids in subclades 1 and 2 (all mosquito-borne viruses) and the 73[K/R]-K-I-[K/R]-[K/R]77 motif of capsids in subclade 4 (all tick-borne viruses) were observed to be completely conserved.

### 3.2 Disorder analysis

The composition of the disorder-promoting amino acids, viz. Ala, Arg, Gly, Gln, Ser, Glu, Asp, His, Lys, and Pro, and the order-promoting amino acids, viz. Ile, Leu, Val, Trp, Tyr, Phe, Thr, Cys, Met, and Asn, were identified and tabulated (**Supplementary Table S2**) (Redwan et al. 2019). The percentage of disorder-promoting amino acids in the C sequences ranged from 45-58%, with the N-terminal region containing a high concentration of disorder-promoting residues (**Table 1**).

The disorder probability plots from PrDOS scores (Ishida and Kinoshita 2007) were compared for all 25 samples to identify conserved patterns. The disorder pattern showed striking similarity within individual subclades (**Fig 2**). From the disorder probability plots, we observed that the N terminal (0-13) and C terminal (110 onwards) residues were disordered across all subclades. In addition, all the subclades showed a peak close to the C-terminus (except the dengue viruses), with viruses in subclade 1 having a notable disorder from residues 94-106. Unlike other viruses, those from subclade 4 showed an extended region of disorder at the N terminus (1-27). Proteins with PPID ≤ 10% are considered highly ordered, those with PPID of 10-30% are moderately disordered, and the ones with PPID ≥ 30% are deemed highly disordered (Redwan et al. 2019). As per this criterion, all the C sequences were found to be moderately disordered (**Table 1**). The ALKV, TBEV, LIV, and OHFV have the highest PPID, whereas the SLEV, ILHV, BANV, DENV2, and DENV4 have the lowest.

The protein binding sites predicted by MoRF_CHiBi_Web_ (Malhis et al. 2016) and DISOPRED3 (Jones and Cozzetto 2015) are presented in (**Table 2**). The motifs observed in the MSA are seen to be a part of the MoRFs of all the capsids, and the residues have undergone only conserved substitutions.

**Table 2:**
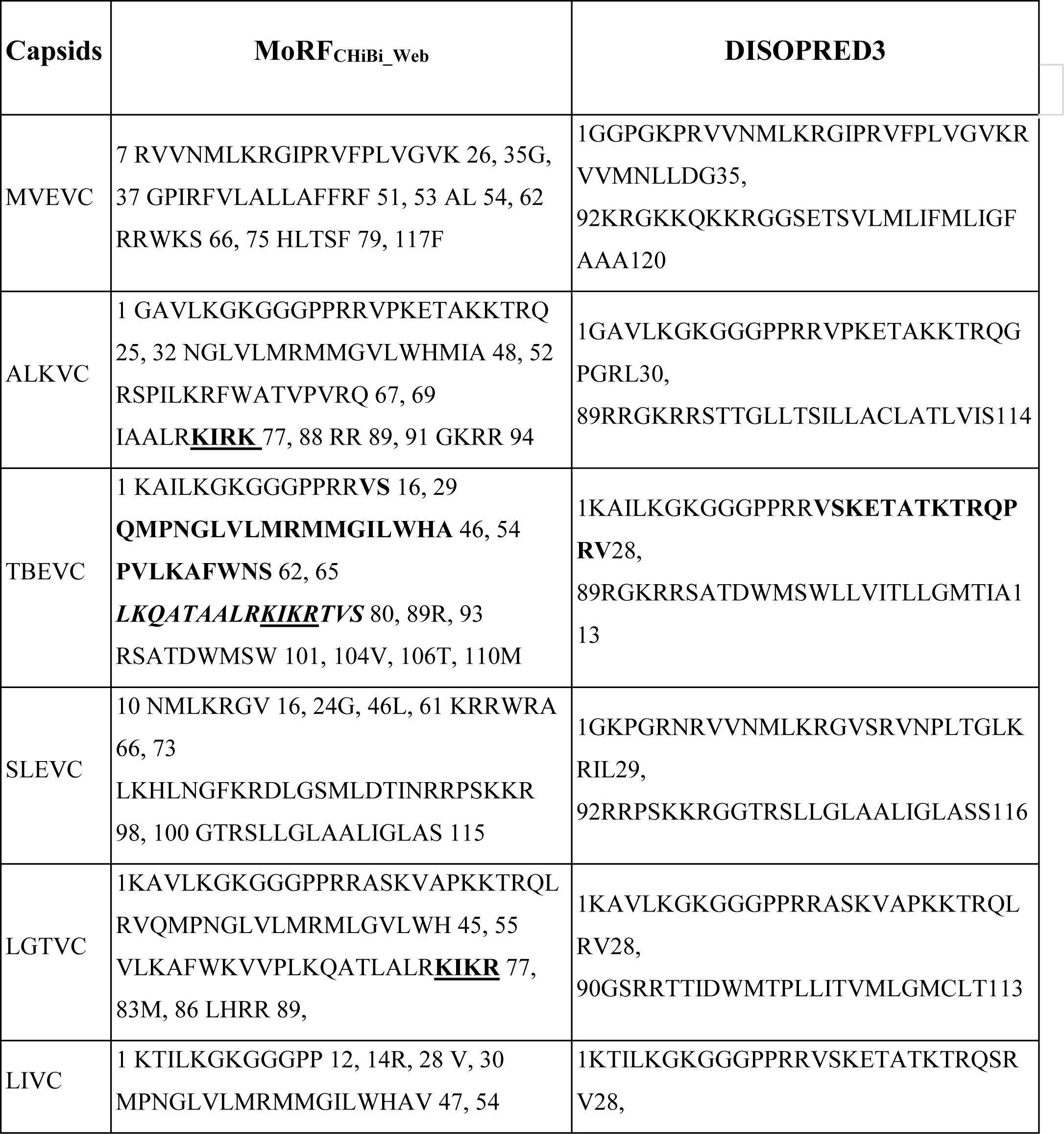
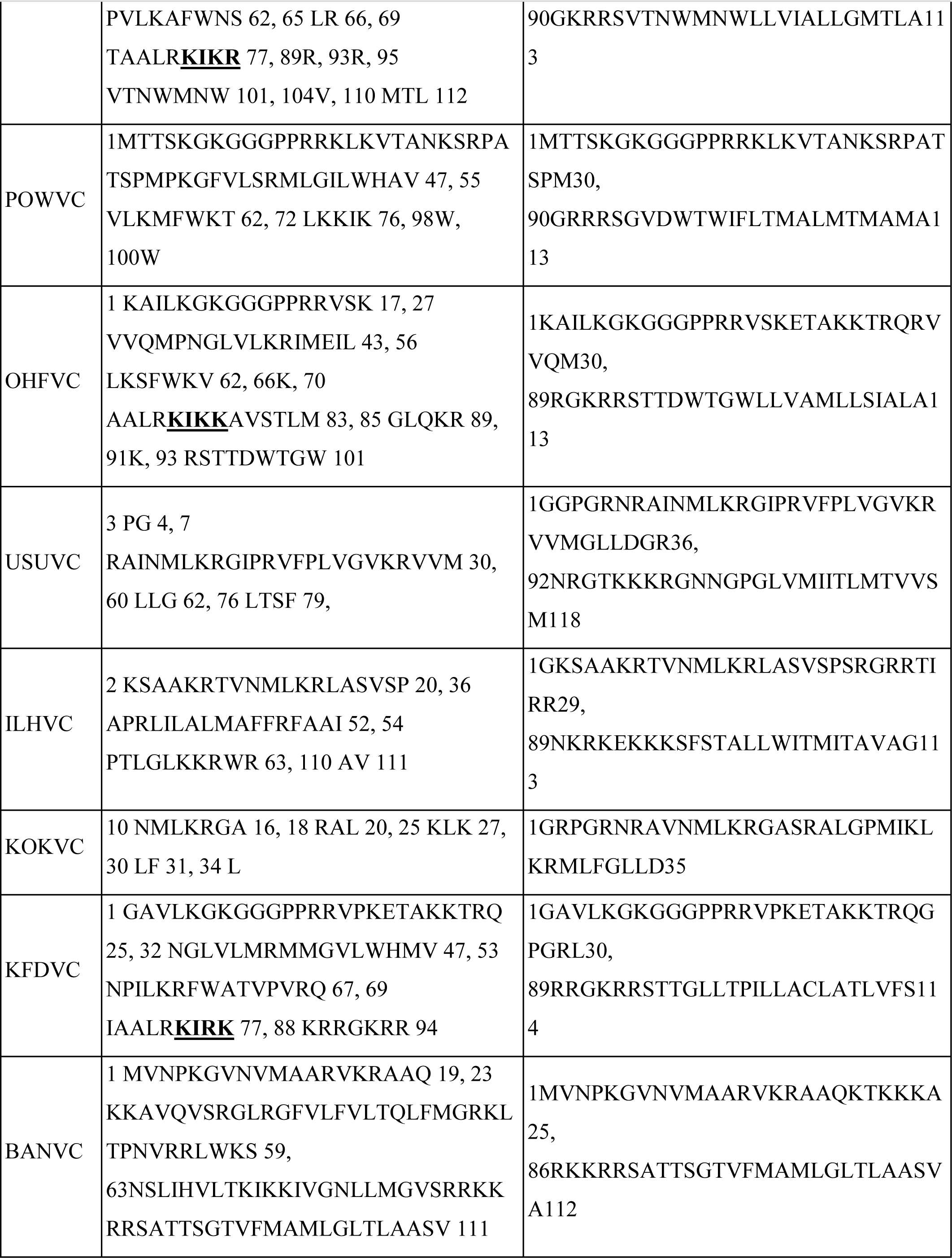
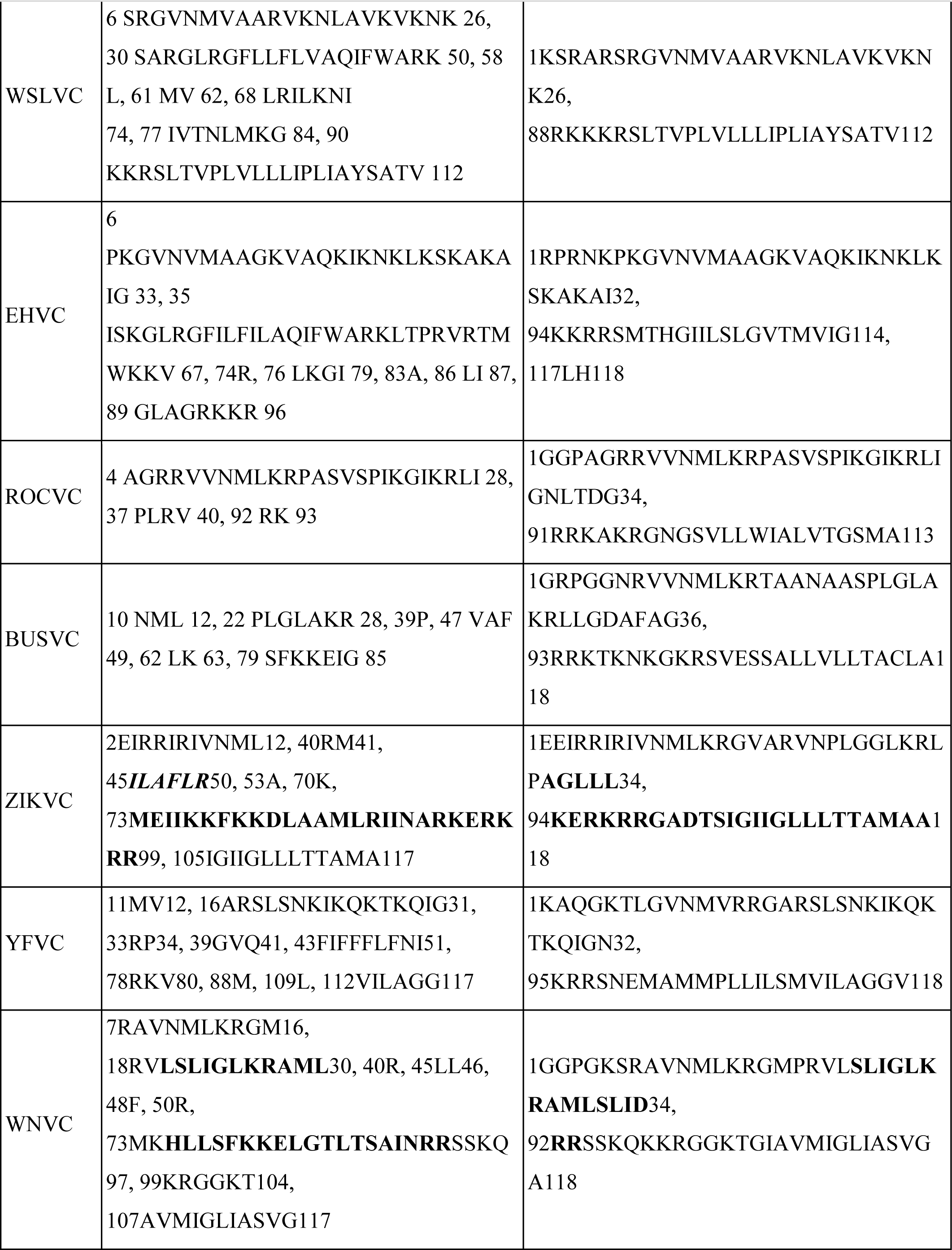
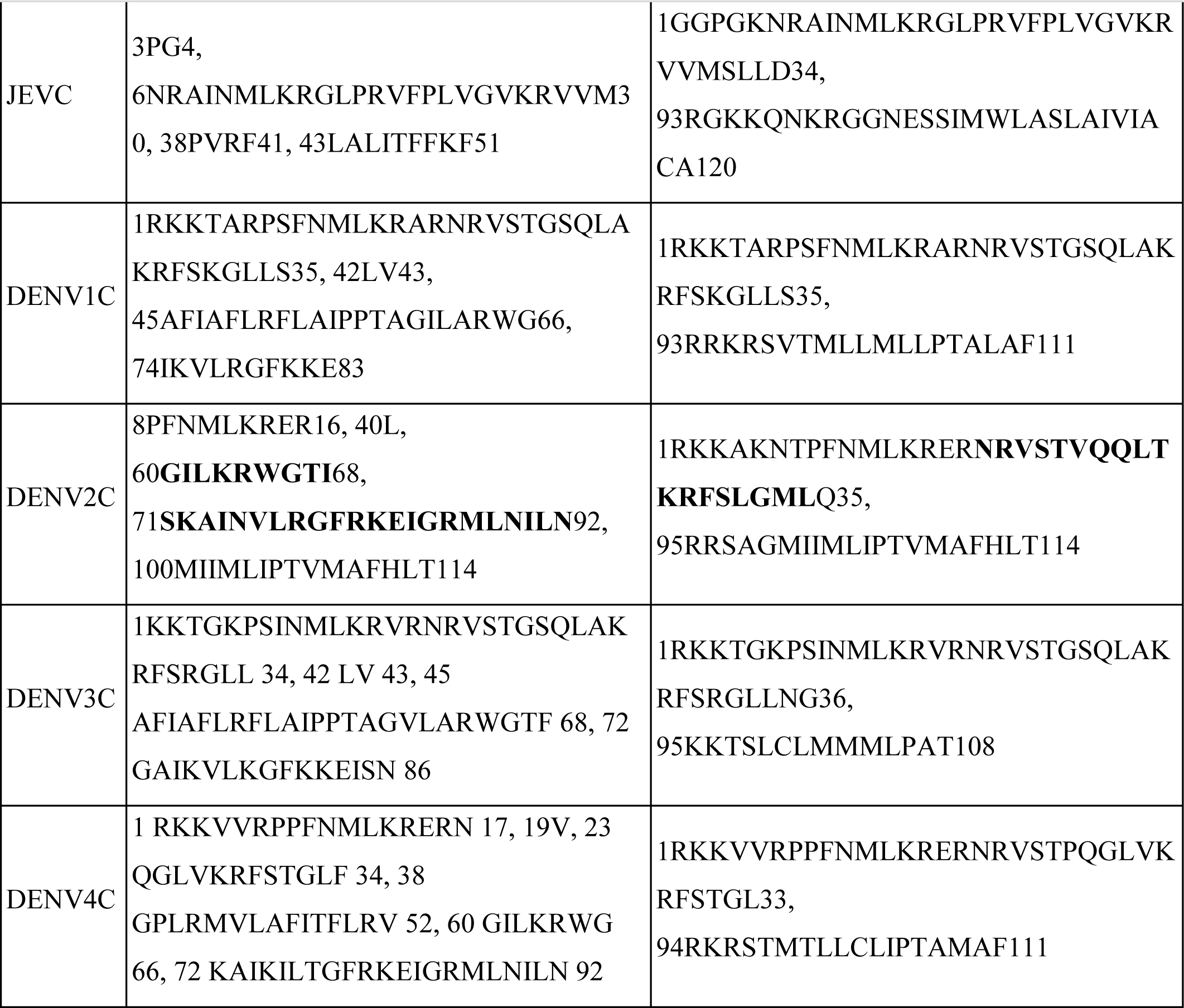
MoRFs predicted by MoRFCHiBi_Web and DISOPRED3. **Boldface-** MoRFs that interact with all the 14 common interacting host proteins of DENVC, ZIKVC, WNVC, and TBEVC. ***Italicized boldface-*** Unique MoRFs that interact with host proteins of similar functions. **<u>Underlined boldface</u>-** Minor binding site of TBEVC NLS with Importin alpha.(Jones and Cozzetto 2015; Malhis et al. 2016)

### 3.3 Interactome analysis & enrichment

From the C-specific interaction network obtained in Cytoscape 3.9.1 (Shannon et al. 2003), the genes encoding common interacting partners for various capsid proteins were identified (as highlighted in **Fig 3**) and listed in **Table 3**.

**Fig 3:**
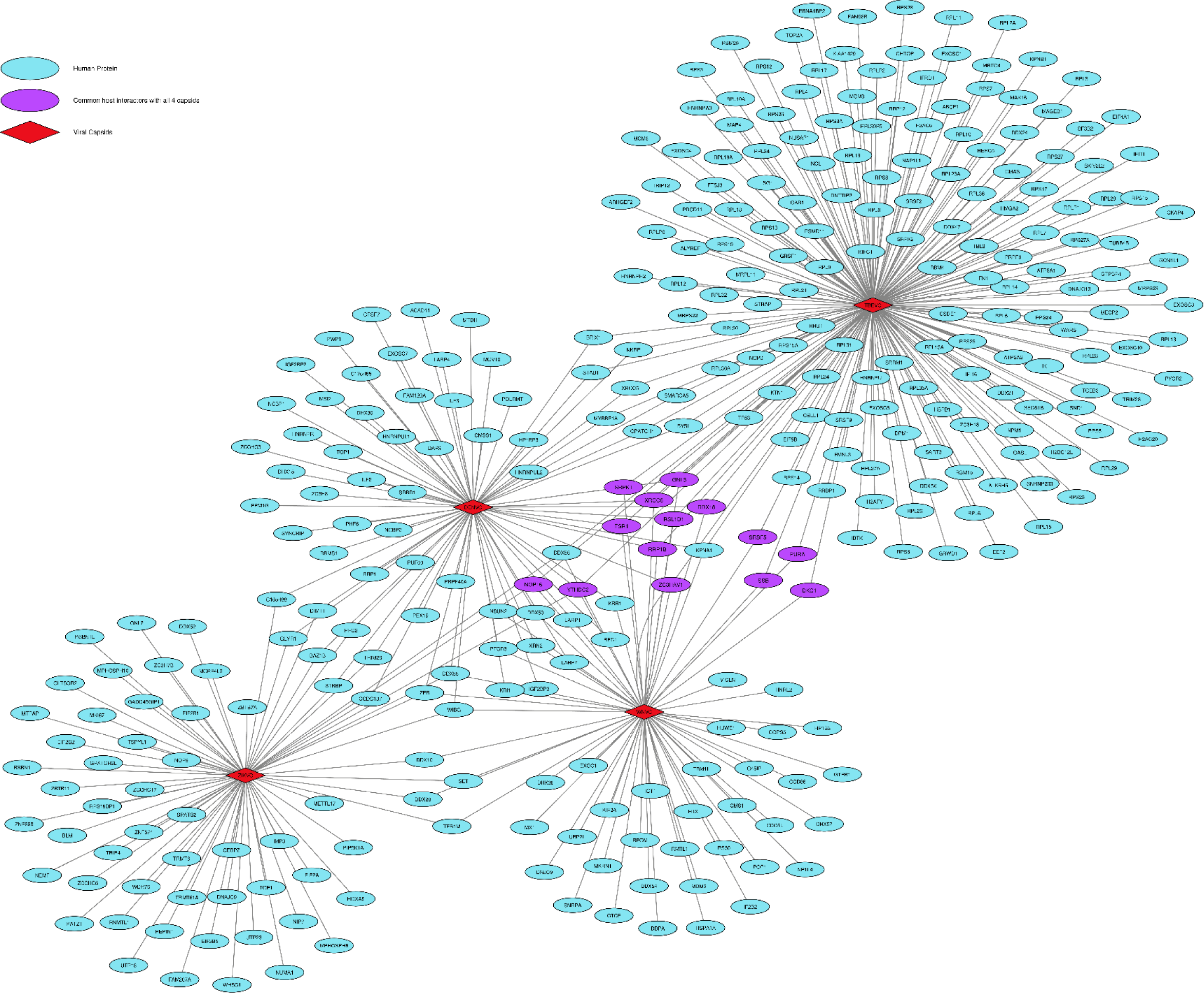
Capsid-Human PPI Network. Representative Network of DENVC, TBEVC, WNVC, and ZIKVC–Human PPI (Li et al. 2019; Selinger et al. 2022).

**Table 3:** Details of Analyzed Human Proteins. Genes encoding common interacting partners of TBEVC, DENV2C, ZIKVC, and WNVC

| Gene Name | Uniprot Id | PDB/Alphafold ID |
| --- | --- | --- |
| TSR1 | Q2NL82 | 6G18 |
| DDX18 | Q9NVP1 | 3LY5 |
| GNL3 | Q9BVP2 | AF-Q9BVP2-F1 |
| RRP1B | Q14684 | AF-Q14684-F1 |
| RSL1D1 | O76021 | AF-O76021-F1 |
| SRPK1 | Q96SB4 | 1WAK |
| ZC3HAV1 | Q7Z2W4 | AF-Q7Z2W4 |
| YTHDC2 | Q9H6S0 | AF-Q9H6S0-F1 |
| DKC1 | O60832 | 7TRC |
| PURA | Q00577 | AF-Q00577-F1 |
| SRSF5 | Q13243 | AF-Q13243-F1 |
| SSB | P05455 | AF-P05455-F1 |
| XRCC6 | P12956 | 1JEY |
| NOP16 | Q9Y3C1 | AF-Q9Y3C1-F1 |

The interactomes of DENV C, ZIKV C, and WNV C were obtained from a study (Li et al. 2019) where affinity-tag purification mass spectrometry (AP-MS) was used to identify the interaction of viral proteins with the host. Similarly, the TBEV C interactome was derived from a study that used co-immunoprecipitation in conjunction with mass spectrometry (Selinger et al. 2022). A total of 1900 interactions were analyzed (**Supplementary Table S3**), and a Venn diagram was constructed based on the human interactors of the four capsids. Fourteen host proteins were identified to be common interactors to all the Cs (**Fig 4**).

**Fig 4:**
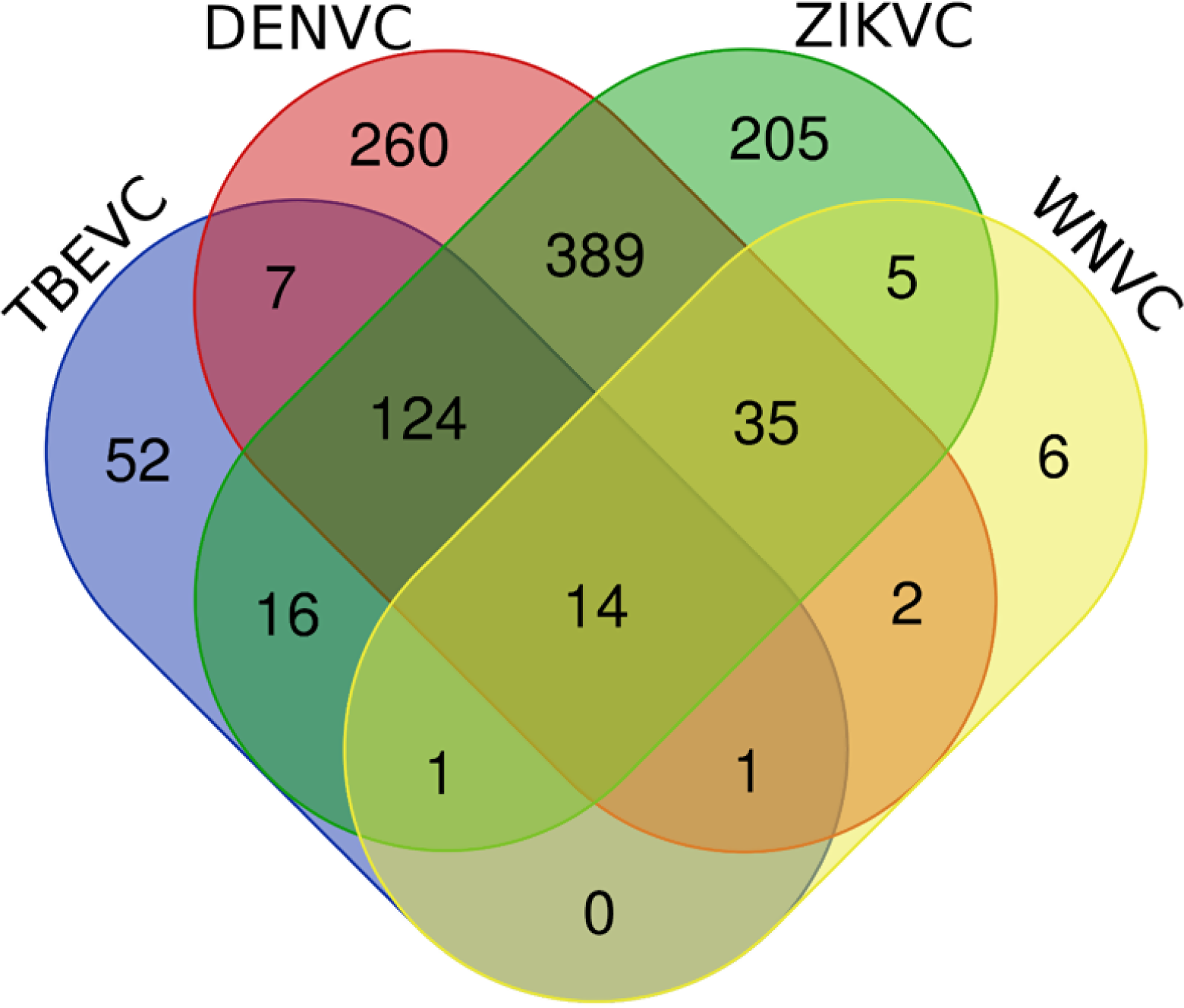
Venn Diagram. Venn diagram showing the number of common interacting host proteins for the selected flaviviruses

Functional enrichment analysis was performed for the common host proteins of all four flaviviruses using g:Profiler (Raudvere et al. 2019). RNA binding and ribosome biogenesis were identified as the most significant molecular function and biological process, respectively. RNA binding mechanism was observed as an enriched function for the shared proteins among Cs of DENV, WNV, and ZIKV C, and between DENV, TBEV, and ZIKV Cs. The cellular component analysis showed that many human proteins interacting with Cs were located in the nucleus, indicating their significance in promoting the replication of viral RNA and gene regulation (**Supplementary Table S5**). It also implies the translocation of the Cs into the nucleus, aided by the protein Importin ɑ, an interacting partner detected in the interactome. DPM1 (a catalytic subunit of the DPMS complex), a human protein shared between DENV C and TBEV C, was reported in a study as an essential host factor to mediate viral replication in DENV and ZIKV infections (Labeau et al. 2020). The biological process, negative regulation of viral genome replication is enriched (IFI16, OASL, HMGA2, and IFIT1) uniquely to TBEV C, which could imply that these host proteins play a role in antiviral defense.

The closely related flaviviruses, DENV C and ZIKV C, have a higher number (389) of commonly interacting host proteins, implying their similar mode of life cycle. Also, the number of MoRF residues observed in the two viruses was quite similar compared to the other two viruses. From the heatmap, it appears that TBEVC and DENVC have a greater number of interacting MoRF residues between the virus-host protein complexes than ZIKV C and WNV C. Notably, TBEV C, the distantly related flavivirus, has a set of 52 unique host-protein interactors. DENV C, ZIKV C, and TBEV C have the second-most common interacting partners of the host (124). Though the phylogenetic analysis shows that C of WNV relates more to the Cs of DENV and ZIKV than TBEV C, the former three viruses have only 35 common interactors, probably owing to limited interaction data (**Fig 4**).

### 3.4 Docking and MoRF Analysis

Protein-protein docking was performed for the flavivirus Cs, TBEV, DENV, ZIKV, and WNV, whose interactome information was previously analyzed. The homodimers of individual Cs were docked against the 14 common human interactors (**Table 3**) using HDOCK (Yan et al. 2017, 2020). The interface residues between the receptor and the ligand (residues within 5.0 Å of each other) were identified from the docking data and compared against the predicted MoRF residues (**Table 2)**. Each C contains a specific MoRF, which interacts with all the docked human proteins (TBEV C: 56-64; DENV C: 75-100; ZIKV C: 78-104; WNV C: 78-96). Additionally, NOP16, SRPK1, SSB, XRCC6, and YTHDC2 interact with the MoRF 67-82 (MCW) of TBEV C, and SRPK1, and ZC3HAV1 interact with residues 50-55 (MCW) of ZIKV C. The percentage of interface residues that are identified to be in MoRFs (**Supplementary Table S4**) can be visualized with the heat map (**Fig 5**).

**Fig 5:**
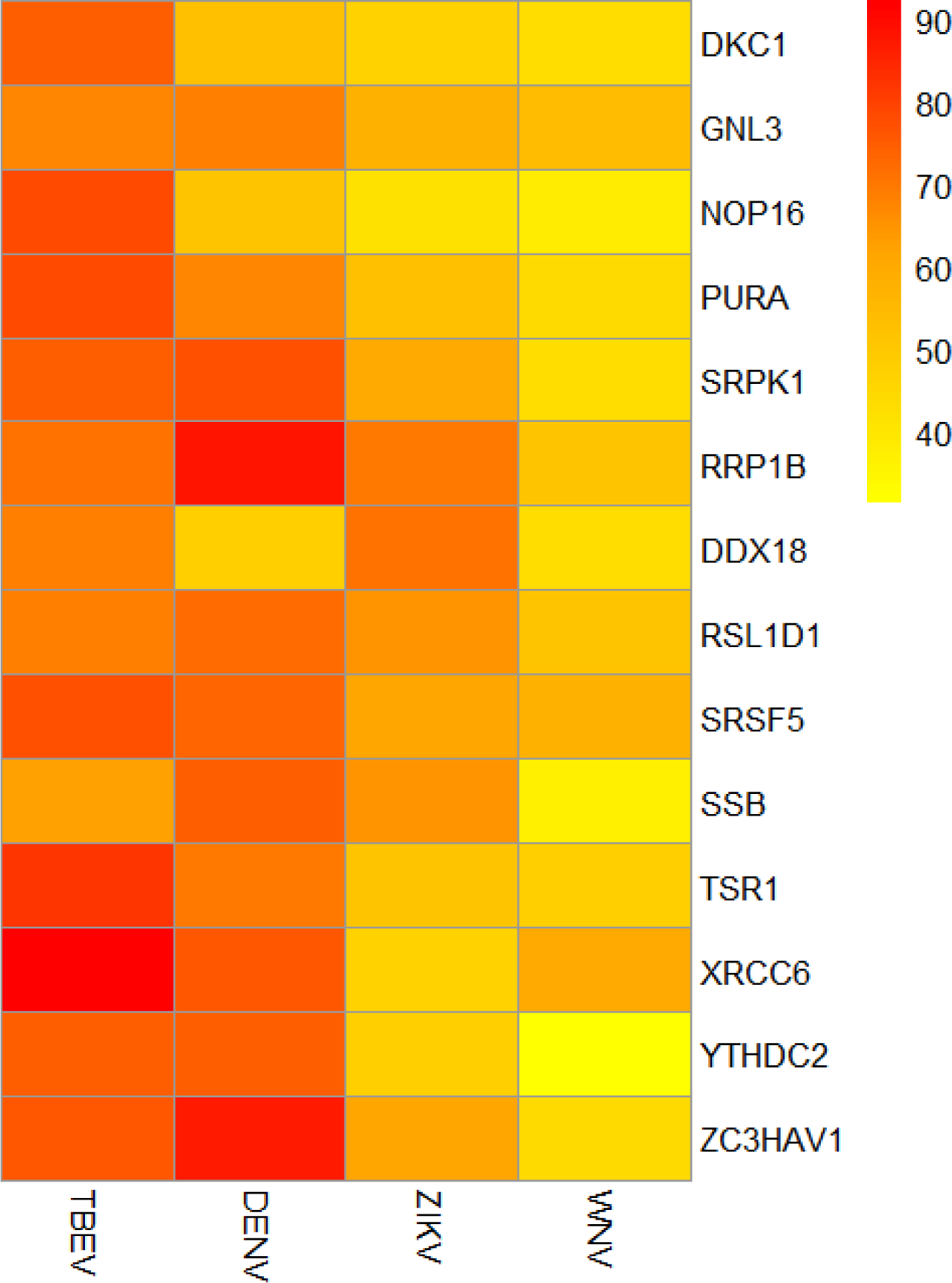
Heatmap. Heatmap representing the percentage of interacting MoRF residues found at the interface of the docked complexes.

## 4. Discussion

The C protein of flaviviruses is found to be an excellent tracker of phylogeny. Changes in the polyprotein and the C, when compared, show a striking similarity, thus making them reliable markers for evolutionary analysis (Martins and Santos 2020). Our results indicate that MoRFs and, by extension, IDRs of flaviviral Cs play an important role in their interactions with multiple host proteins. Furthermore, it was observed that the underlying disorder had been conserved in each subclade. Certain MoRFs, which were identified across the subclades interacted with the same host proteins, proving that the interactions are indispensable during flaviviral infections.

The Cs of mosquito-borne flaviviruses have significantly overlapping interactomes, implying functional and structural conservation despite low sequence similarity. For example, DENV and ZIKV show remarkable similarities in their transmission, replication cycle, and disease manifestations like congenital microcephaly and developmental defects in infants (Zhang et al. 2021). Hence, comparative analysis of their interactomes aided in identifying integral proteins that could be potential targets for antiviral therapies. The DENV and ZIKV Cs interact with multiple proteins involved in RNA processing, transcription, and DNA replication, which mandates the translocation of C to the nucleus (Shah et al. 2018). DENV C MoRF has an importin-α like motif, which might facilitate nuclear entry via the importin α/β complex. Similar motifs are found in several mosquito-borne flaviviruses suggesting a common scheme of nuclear transport (Martins and Santos 2020). Surprisingly, the MoRFs of ALKV, TBEV, LGTV, LIV, OHFV, and KFDV, which are transmitted by ticks, also feature similar MoRFs. The binding sequence at the minor site, 74KIKR77 (shown as **<u>KIKR</u>** in **Table 2)**, is highly conserved in these viruses, and it was positioned at the interface of TBEVC NLS (nuclear localization signal) and importin α in the co-crystal structure (Selinger et al. 2022). Studies with ALKV also corroborate the role of MoRFs in such interactions (Redwan et al. 2019). Comparative screening of the polyprotein interactome of different flaviviruses, helps us identify host interactors specific for Cs. For example, TDIF2 (that regulates the transcriptional activity and chromatin remodelling) interacts with the Cs of TBEV via its MoRF (Selinger et al. 2022). Hence, the presence of the conserved MoRFs within the TBEV subclade suggests the possibility of similar interactions of other viruses with TDIF2.

According to the heatmap of interactions based on docking analyses of 14 common host proteins with Cs of TBEV, DENV, ZIKV, and WNV, the former two have the highest percentage of interacting MoRF residues. ZIKV and WNV, classified under Subclade 1 in the phylogenetic analysis with PPIDs of 14.41% and 16.95%, respectively, show fewer interacting MoRF residues. A higher PPID (26.55%) observed for TBEV explains the increased prevalence of MoRFs at the interface in many common host proteins (TBEV binding). Following TBEV, DENV2, with a PPID of 12.28%, also shows a higher number of interacting MoRF residues for most of the host proteins. Other studies have established a similar correlation between the degree of disorder and virulency in different viruses (Goh et al. 2015, 2016). TSR1, one of the common human interactors identified, was found to be involved in the binding of viral proteins leading to a translational shutdown in SARS-COV2-infected cells (Thoms et al. 2020). Thus, IDRs might be useful as a drug target as it was suggested that a high disorder level in the C influenced SARS-CoV-2 virulence (Tenchov and Zhou 2022). Our study reveals that regardless of the level of disorder in the Cs, interactions with MoRFs are mandatory for the establishment of a successful infection (MoRFs shown in **boldface** in **Table 2**).

The fact that the Cs of ZIKV, DENV, WNV, and TBEV had phylogenetically diverged to utilize different vectors, infect different hosts, and showed varied manifestations of the infection, and yet interacted via their MoRFs with 14 host proteins, emphasizes the importance of order in the disordered regions. However, the MoRFs were also observed to interact with unique host proteins. NOP16, SRPK1, SSB, XRCC6, and YTHDC2 interact with the MoRF 67-82 (MCW) (***Italicized boldface*** in **Table 2)** of TBEV C. Their functional enrichment reveals involvement in mRNA processing and maturation, ribosome biogenesis, and RNA binding. It also interacts with the protein kinase domain (SRPK1) and YT521-B-like domain (YTHDC2), both of which modulate alternate splicing. Similarly, SRPK1 and ZC3HAV1 interact with residues 50-55 (MCW) (***Italicized boldface*** in red in **Table 2)** of ZIKVC. Both proteins have antiviral functions such as decreasing viral genome packaging and viral mRNA degradation, respectively. The Cs of ZIKV, DENV, and WNV are known to interact with the exon-junction complex (EJC) recycling factor, PYM1, which has an anti-viral function and influences nonsense-mediated decay of viral RNA. The interaction of C with PYM1 leads to its sequestration, inhibiting anti-viral activity and protecting viral RNA from degradation (Li et al. 2019). ZC3HAV1 is also involved in antiviral immune response as it interacts with IFIX, which recognizes viral DNA in the nucleus (Diner et al. 2015). Thus, the structural flexibility of MORFs has facilitated their utilization for varied interactions during pathogenesis.

## 5. Conclusion

Through this analysis, we detected conserved patterns of IDRs within Cs of the same subclades and observed that the capsid interactions with human proteins depend on residues present in the MoRF regions. The capsid interactions are conserved in flaviviruses, as viruses from different subclades have common interactors despite differences in sequence. The extensive interactomes revealed that each MoRF in the flaviviral C binds to multiple partners responsible for vital functions like immune response, gene expression, transcriptional and translational regulation. The study thus provides evidence supporting the role of IDRs of Cs in the evolution and virulence of the flaviviruses. These results highlight the importance of IDRs as a gateway for new strategies against viral infections and the development of novel therapeutics.

## ACKNOWLEDGEMENT

SV thanks University Grants Commission-Faculty Recharge Program (UGC-FRP), New Delhi, India, for salary support. The authors thank BIC at DoBT, AU (BT/PR40163/BTIS/137/31/2021), DBT, Govt. of India for computational facilities.

## AUTHOR CONTRIBUTION

AS, PS and PU conceptualized the idea, collected the data, carried out the analysis, and wrote the paper. KR was involved in interaction and docking studies and drafted the corresponding sections. SV was involved in design of the study, analysis, and review of the manuscript. All authors have reviewed the manuscript before submission.

## CONFLICT OF INTEREST/COMPETING INTEREST

On behalf of all authors, the corresponding author states that there is no conflict of interest.

## FUNDING

No funding was received for conducting this study.

## DATA AVAILABILITY

The data that support the findings of this study are provided in the supplementary information.

## DECLARATIONS

Dr. Sangita Venkataraman receives her salary from University Grants Commission-Faculty Recharge Program (UGC-FRP), New Delhi. No funding was received for conducting the study.

## CONSENT

Informed consent was obtained from all individual participants included in the study.

## Notes

### Competing Interest Statement

The authors have declared no competing interest.

## REFERENCES

Abor Erd G′, Os ″, Atyásatýatyás Pajkos M′, et al (2021) IUPred3: prediction of protein disorder enhanced with unambiguous experimental annotation and visualization of evolutionary conservation. Nucleic Acids Res 49:W297–W303. 10.1093/NAR/GKAB408

Barik S (2020) Genus-specific pattern of intrinsically disordered central regions in the nucleocapsid protein of coronaviruses. Comput Struct Biotechnol J 18:1884–1890. 10.1016/j.csbj.2020.07.005

Bateman A, Martin MJ, Orchard S, et al (2021) UniProt: the universal protein knowledgebase in 2021. Nucleic Acids Res 49:D480–D489. 10.1093/NAR/GKAA1100

Berman HM (2000) The Protein Data Bank. Nucleic Acids Res 28:235–242. 10.1093/nar/28.1.235

Byk LA, Gamarnik A V. (2016) Properties and Functions of the Dengue Virus Capsid Protein. Annu Rev Virol 3:263–281. 10.1146/annurev-virology-110615-042334

De Madrid AT, Porterfield JS (1974) The flaviviruses (group B arboviruses): a cross neutralization study. Journal of General Virology 23:91–96. 10.1099/0022-1317-23-1-91/CITE/REFWORKS

Diner BA, Li T, Greco TM, et al (2015) The functional interactome of PYHIN immune regulators reveals IFIX is a sensor of viral DNA. Mol Syst Biol 11:787. 10.15252/msb.20145808

Faustino, Martins, Karguth, et al (2019) Structural and Functional Properties of the Capsid Protein of Dengue and Related Flavivirus. Int J Mol Sci 20:3870. 10.3390/ijms20163870

Goh G, Dunker A, Foster J, Uversky V (2019) Zika and Flavivirus Shell Disorder: Virulence and Fetal Morbidity. Biomolecules 9:710. 10.3390/biom9110710

Goh GK-M, Dunker AK, Uversky VN (2016) Correlating Flavivirus virulence and levels of intrinsic disorder in shell proteins: protective roles vs. immune evasion. Mol Biosyst 12:1881–1891. 10.1039/C6MB00228E

Goh GK-M, Dunker AK, Uversky VN (2015) Detection of links between Ebola nucleocapsid and virulence using disorder analysis. Mol Biosyst 11:2337–2344. 10.1039/C5MB00240K

Ishida T, Kinoshita K (2007) PrDOS: prediction of disordered protein regions from amino acid sequence. Nucleic Acids Res 35:. 10.1093/NAR/GKM363

Ivanyi-Nagy R, Lavergne J-P, Gabus C, et al (2008) RNA chaperoning and intrinsic disorder in the core proteins of Flaviviridae. Nucleic Acids Res 36:712–725. 10.1093/nar/gkm1051

Jones DT, Cozzetto D (2015) DISOPRED3: precise disordered region predictions with annotated protein-binding activity. Bioinformatics 31:857. 10.1093/BIOINFORMATICS/BTU744

Jumper J, Evans R, Pritzel A, et al (2021) Highly accurate protein structure prediction with AlphaFold. Nature 596:583–589. 10.1038/s41586-021-03819-2

Labeau A, Simon-Loriere E, Hafirassou M-L, et al (2020) A Genome-Wide CRISPR-Cas9 Screen Identifies the Dolichol-Phosphate Mannose Synthase Complex as a Host Dependency Factor for Dengue Virus Infection. J Virol 94:. 10.1128/JVI.01751-19

Li M, Johnson JR, Truong B, et al (2019) Identification of antiviral roles for the exon–junction complex and nonsense-mediated decay in flaviviral infection. Nat Microbiol 4:985–995. 10.1038/s41564-019-0375-z

Malhis N, Jacobson M, Gsponer J (2016) MoRFchibi SYSTEM: software tools for the identification of MoRFs in protein sequences. Nucleic Acids Res 44:W488–W493. 10.1093/nar/gkw409

Martins IC, Santos NC (2020) Intrinsically disordered protein domains in flavivirus infection. Arch Biochem Biophys 683:108298. 10.1016/j.abb.2020.108298

Mishra PM, Verma NC, Rao C, et al (2020) Intrinsically disordered proteins of viruses: Involvement in the mechanism of cell regulation and pathogenesis. In: Progress in Molecular Biology and Translational Science. Elsevier B.V., pp 1–78

Neves V, Aires-da-Silva F, Morais M, et al (2017) Novel Peptides Derived from Dengue Virus Capsid Protein Translocate Reversibly the Blood–Brain Barrier through a Receptor-Free Mechanism. ACS Chem Biol 12:1257– 1268. 10.1021/acschembio.7b00087

Paysan-Lafosse T, Blum M, Chuguransky S, et al (2023) InterPro in 2022. Nucleic Acids Res 51:D418–D427. 10.1093/nar/gkac993

Quaglia F, Mészáros B, Salladini E, et al (2022) DisProt in 2022: improved quality and accessibility of protein intrinsic disorder annotation. Nucleic Acids Res 50:D480–D487. 10.1093/nar/gkab1082

Raudvere U, Kolberg L, Kuzmin I, et al (2019) g:Profiler: a web server for functional enrichment analysis and conversions of gene lists (2019 update). Nucleic Acids Res 47:W191–W198. 10.1093/nar/gkz369

Redwan EM, AlJaddawi AA, Uversky VN (2019) Structural disorder in the proteome and interactome of Alkhurma virus (ALKV). Cellular and Molecular Life Sciences 76:577–608. 10.1007/s00018-018-2968-8

RStudio Team RStudio version 4.2.2

Selinger M, Novotný R, Sýs J, et al (2022) Tick-borne encephalitis virus capsid protein induces translational shutoff as revealed by its structural-biological analysis. J Biol Chem 298:102585. 10.1016/J.JBC.2022.102585

Shah PS, Link N, Jang GM, et al (2018) Comparative Flavivirus-Host Protein Interaction Mapping Reveals Mechanisms of Dengue and Zika Virus Pathogenesis. Cell 175:1931–1945.e18. 10.1016/j.cell.2018.11.028

Shannon P, Markiel A, Ozier O, et al (2003) Cytoscape: a software environment for integrated models of biomolecular interaction networks. Genome Res 13:2498–2504. 10.1101/GR.1239303

Sharma NR, Gadhave K, Kumar P, et al (2021) Analysis of the dark proteome of Chandipura virus reveals maximum propensity for intrinsic disorder in phosphoprotein. Sci Rep 11:13253. 10.1038/s41598-021-92581-6

Tamura K, Stecher G, Kumar S (2021) MEGA11: Molecular Evolutionary Genetics Analysis Version 11. Mol Biol Evol 38:3022–3027. 10.1093/molbev/msab120

Tenchov R, Zhou QA (2022) Intrinsically Disordered Proteins: Perspective on COVID-19 Infection and Drug Discovery. ACS Infect Dis 8:422–432. 10.1021/acsinfecdis.2c00031

Thoms M, Buschauer R, Ameismeier M, et al (2020) Structural basis for translational shutdown and immune evasion by the Nsp1 protein of SARS-CoV-2. Science (1979) 369:1249–1255. 10.1126/science.abc8665

Varadi M, Anyango S, Deshpande M, et al (2022) AlphaFold Protein Structure Database: massively expanding the structural coverage of protein-sequence space with high-accuracy models. Nucleic Acids Res 50:D439–D444. 10.1093/nar/gkab1061

Wubben JM, Atkinson SC, Borg NA (2020) The Role of Protein Disorder in Nuclear Transport and in Its Subversion by Viruses. Cells 9:2654. 10.3390/cells9122654

Yan Y, Tao H, He J, Huang SY (2020) The HDOCK server for integrated protein–protein docking. Nature Protocols 2020 15:5 15:1829–1852. 10.1038/s41596-020-0312-x

Yan Y, Zhang D, Zhou P, et al (2017) HDOCK: a web server for protein-protein and protein-DNA/RNA docking based on a hybrid strategy. Nucleic Acids Res 45:W365–W373. 10.1093/NAR/GKX407

Yves Van de Peer Bioinformatics & Evolutionary Genomics. http://bioinformatics.psb.ugent.be/people. Accessed 16 Apr 2023

Zhang X, Zhang Y, Jia R, et al (2021) Structure and function of capsid protein in flavivirus infection and its applications in the development of vaccines and therapeutics. Vet Res 52:98. 10.1186/s13567-021-00966-2

